# Bi-directional Mendelian randomization of epithelial ovarian cancer and schizophrenia and uni-directional Mendelian randomization of schizophrenia on circulating glycerophosphocholine metabolites

**DOI:** 10.1101/655894

**Authors:** Charleen D. Adams, Susan L. Neuhausen

**Affiliations:** Department of Population Sciences, Beckman Research Institute of City of Hope, Duarte, CA, USA

**Keywords:** Schizophrenia, ovarian cancer, Mendelian randomization, glycerophosphocholine, metabolism

## Abstract

Most women with epithelial ovarian cancer (EOC) present with late-stage disease. As a result, globally, EOC is responsible for more than 150,000 deaths a year. Thus, a better understanding of risk factors for developing EOC is crucial for earlier screening and detection to improve survival. To that effort, there have been suggestions that there is an association of schizophrenia and cancer, possibly because metabolic changes are a hallmark of both cancer and schizophrenia (SZ). Perturbed choline metabolism has been documented in both diseases. Our objective was to use Mendelian randomization to evaluate whether SZ increased risk for developing EOC or the converse, and, whether SZ impacted glycerophosphocholine (GPC) metabolites. We found that SZ conferred a weak but increased risk for EOC, but not the reverse (no evidence that EOC caused SZ). SZ was also causally associated with lower levels of two GPC species and with suggestively lower levels in an additional three GPCs. We postulate that perturbed choline metabolism in SZ may mimic or contribute to a “cholinic” phenotype, as observed in EOC cells.

## INTRODUCTION

Though epithelial ovarian cancer (EOC) is rare, across the globe, over 200,000 new cases of ovarian cancer are diagnosed yearly [1]. Because the majority of diagnoses are advanced-stage disease with poor prognosis and cancer recurrence after treatment [2],[3], globally, over 150,000 women die from this cancer each year [4]. Aside from family history of breast or ovarian cancer, high body mass index in adolescence, diagnosis of endometriosis, and taller height, most established risk factors are hormonal or reproductive [1]. A recent Mendelian randomization (MR) study of 13 known observational risk factors only confirmed genetic liability to lifetime smoking and endometriosis as causal [5]. Thus, identification of additional risk factors is crucial for designing prevention strategies and for earlier diagnosis of EOC.

Because EOC and breast cancer share many similar risk factors and schizophrenia (SZ) has been reported to be a causal risk factor for breast cancer in a recent Mendelian randomization (MR) study [6], we hypothesized that SZ or something about its disease pathogenesis, perhaps metabolic, may be a risk factor for EOC. One possibility is energy metabolism, as it is important in both cancer [7] and schizophrenia (SZ) [8], [9]. In particular, aberrant choline metabolism has been documented for EOC [10], [11] and also noted in the brains of schizophrenics [8]. We conducted an MR study of SZ and EOC and its subtypes, and, to assess whether EOC might influence SZ, we also tested the reverse: the causal impact of EOC on SZ. As proof of principle that SZ may impact on circulating metabolites that have been associated with EOC, we performed MR of SZ on 15 circulating glycerophosphocholine (GPC) metabolites.

## MATERIALS AND METHODS

### Conceptual framework

MR is an instrumental-variables technique that exploits the random assortment of alleles, using, most commonly, single-nucleotide polymorphisms (SNPs) as instruments (“proxies”) [12], [13]. With summary results from genome-wide association studies (GWASs), we used two-sample MR to test the causal relationships between 1) EOC and SZ and 2) SZ and GPCs. We presumed the following assumptions: the chosen instrumental variables (SNPs) are strongly associated with exposures (assumption 1); the instrumental variables are not associated with confounders (assumption 2); and the instrumental variables are only associated with the outcomes through the exposures (assumption 3) [14].

### Data Sources

Two-sample MR uses the summary results from two separate GWASs as its data sources: a GWAS of the exposure and a GWAS of the outcome.

#### SZ on EOC and EOC subtypes

1. We selected a GWAS of SZ [15]; Ripke *et al.* (2014) meta-analyzed 49 case-control studies of SZ along with 1,235 parent affected-offspring trios (total study size = 35,476 SZ cases and 46,839 controls). From their GWAS results, we extracted the summary statistics for SNPs strongly associated with SZ (p-value <5e-8).
2. Next, we identified the largest GWAS of EOC and its subtypes [16]; Phelan *et al.* (2017) meta-analyzed GWAS from six Ovarian Cancer Association Consortium (OCAC) and two Consortium of Investigators of Modifiers of BRCA1/2 (CIMBA) projects (total study size = 25,509 cases and 40,941 controls of European ancestry) for women who had been genotyped using the Illumina Custom Infinium array (OncoArray). We extracted summary data for the SZ-associated SNPs from within the GWAS of overall EOC (n=25,509), high-grade serous carcinoma (n=13,037), mucinous carcinoma (n=2,566), endometrioid carcinoma (n=2,810), and clear-cell carcinoma (n=1,366).

#### EOC on SZ (reverse direction)

For the MR of EOC on SZ, we selected SNPs strongly associated with EOC (p-value <5e-8) from the Phelan *et al.* (2017) GWAS of EOC.

We extracted the summary data for the EOC-associated SNPs from the Ripke *et al.* (2014) GWAS of SZ.

#### SZ on 15 GPCs

For the MR of SZ on GPCs, we extracted SZ-associated SNPs (p-value <5e-8) from Ripke *et al.* (2014).

We then selected GWASs of circulating metabolites in, depending on the metabolite, up to 7,824 adult individuals from two European population studies (TwinsUK and the Cooperative Health Research in the Region of Augsburg (KORA) study) performed by Shin *et al.* (2014). We extracted summary statistics for SZ-associated SNPs for 15 GPC metabolites: 1-arachidonoylglycerophosphocholine, 1-palmitoleoylglycerophosphocholine, 1-eicosatrienoylglycerophosphocholine, 1-docosahexaenoylglycerophosphocholine, 1-eicosadienoylglycerophosphocholine, 1-palmitoylglycerophosphocholine, 1-heptadecanoylglycerophosphocholine, 1-oleoylglycerophosphocholine, 1-stearoylglycerophosphocholine, 1-linoleoylglycerophosphocholine, 2-palmitoylglycerophosphocholine, 2-oleoylglycerophosphocholine, 2-stearoylglycerophosphocholine, 2-linoleoylglycerophosphocholine, 1-myristoylglycerophosphocholine [17].

### Statistical procedure

#### SZ on EOC and EOC subtypes

To implement MR for the tests of SZ on overall EOC and SZ on EOC subtypes, we used the summary statistics for SZ-associated SNPs identified in and extracted from Ripke *et al.* (2014) and the summary statistics for the Ripke-SZ-associated SNPs extracted from Phelan *et al*. (2017) to calculate the odds ratio (OR) for overall EOC and for each subtype per unit log odds higher liability to SZ for each SNP instrument (Wald ratio) and conducted a meta-analysis of the Wald ratios in a combined instrument with the inverse-weighted variance (IVW) MR method [18].

#### EOC on SZ

To implement MR for the test of EOC on SZ, we used the summary statistics for EOC-associated SNPs identified in and extracted from Phelan *et al.* (2017) and the summary statistics for the Phelan-EOC-associated SNPs extracted from Ripke *et al.* (2014) to calculate the odds ratio (OR) for SZ per unit log odds higher liability to EOC for each SNP instrument (Wald ratio) and meta-analyzed the results with the IVW method. The MR of SZ on EOC and the reverse MR of EOC on SZ comprise a bidirectional analysis. Because the validity of bidirectional MR methods depends on the independence of the two instruments [12], we checked that there was no overlap or linkage disequilibrium (LD) between the genetic variants included in the instruments for SZ and EOC. We used LDassoc, a free and publicly available web tool, to assess whether instruments were in LD [19].

#### SZ on 15 GPCs

To implement MR for the tests of SZ on 15 GPC metabolites, we used the summary statistics for the SZ-associated SNPs identified in and extracted from Ripke *et al.* (2014) and Ripke-SZ-associated SNPs extracted from the Shin *et al.* (2014) GWAS of circulating GPC metabolites. We calculated the log10 increase in GPC metabolite per unit log odds higher liability to SZ for each SNP proxy and meta-analyzed the Wald ratios with the IVW method. We performed 15 meta-analyses, one for each of the 15 GPC metabolites. To account for multiple testing, we set the *P*-value threshold for the analyses of SZ on GPCs to the Bonferroni-corrected value of 0.0033 (*P*=0.05/15).

### Two-Sample MR

For all instruments, where the exposure SNPs were not available in the outcome data, we identified LD proxy SNPs in linkage disequilibrium (R^2^ > 0.8 and within 250kb of the target SNP). We implemented all two-sample MR tests in R version 3.5.1 using the “TwoSampleMR” package [20]. All data are publicly available and accessible through the MR-Base website, a curated database of complete GWASs and platform for performing two-sample MR [21].

### Sensitivity Analyses

To detect possible violations of the third MR assumption (assumption 3 above), we performed various sensitivity analyses. We applied IVW radial regression for each MR test. IVW radial regression uses the *P*-value from Cochran’s *Q*-statistic (*P*[het]) to identify possible heterogeneity among an instrument’s ratio estimates [22]. Heterogeneity can indicate pleiotropy. For all analyses, we removed heterogenous SNPs and report the results of the MR analysis with them removed, along with IVW radial regression performed again after their removal to demonstrate either absence of evidence for heterogeneity or continued evidence for it. The tables of results include the number of SNPs included in each analysis. We also implemented MR-Egger regression, weighted median, and weighted mode estimators, which make different assumptions about the underlying nature of potential pleiotropy. Briefly, if the MR-Egger, weighted median, and weighted mode estimators have the same direction and comparable magnitude of effects as the IVW estimate, this strengthens confidence in the validity of the instruments.

## RESULTS

### SZ and EOC

SZ was associated with overall EOC (IVW OR per log odds higher liability to SZ: 1.09, 95% CI: 1.04, 1.14; *P*=0.0004) and high-grade serous ovarian carcinoma (IVW OR 1.07, 95% CI: 1.01, 1.13; *P*=0.033). The comparisons of the MR estimators for both reveal little evidence for pleiotropy: the direction of the effect was consistent across methods and the effects were of relatively the same magnitude for the IVW, weighted median, and weighted mode estimators (Table 1; Figure 1). IVW radial regression indicated no evidence of heterogeneity (overall EOC: *P*[het]=0.5238; high-grade serous: *P*[het]=0.4254).

**Table 1.**
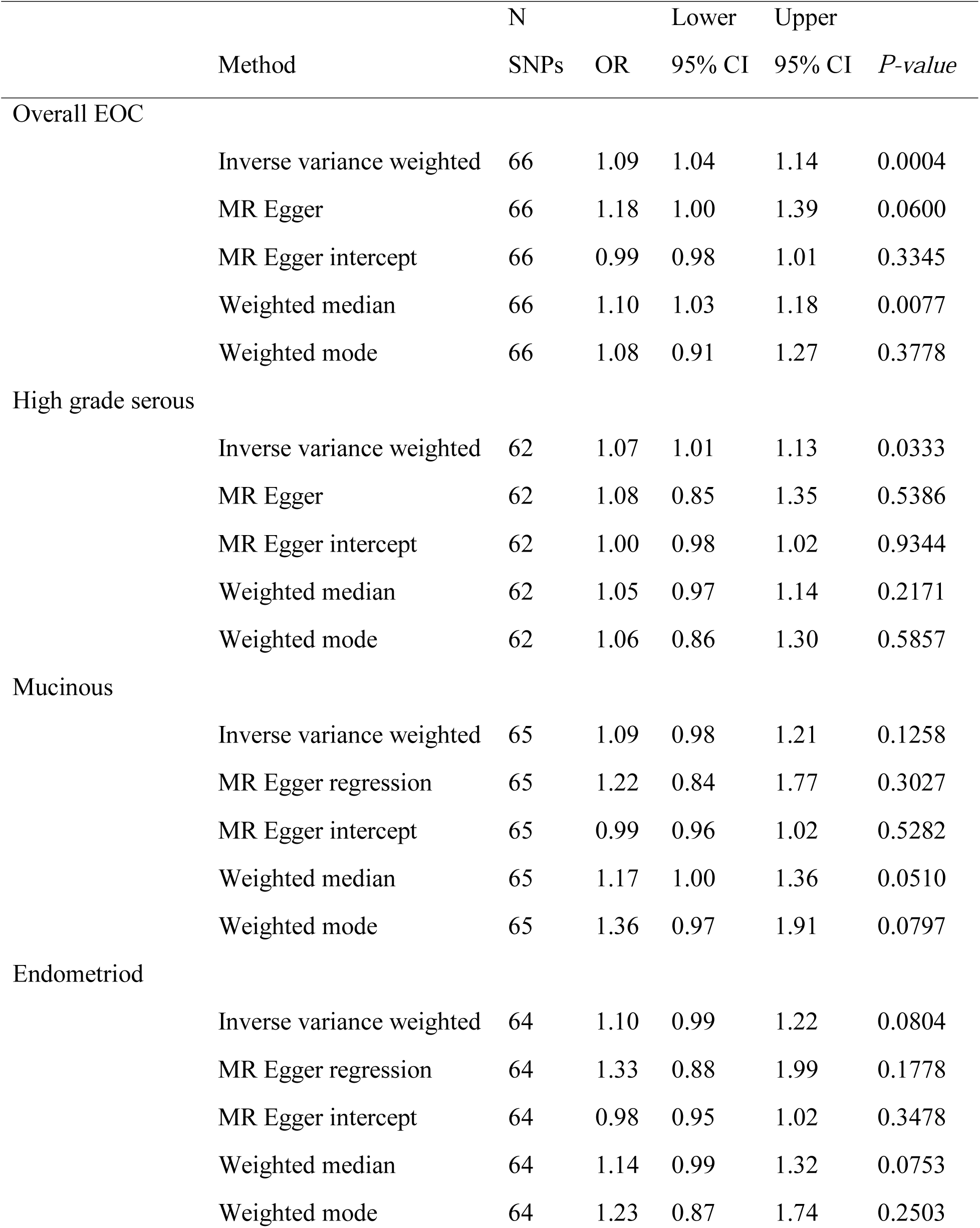
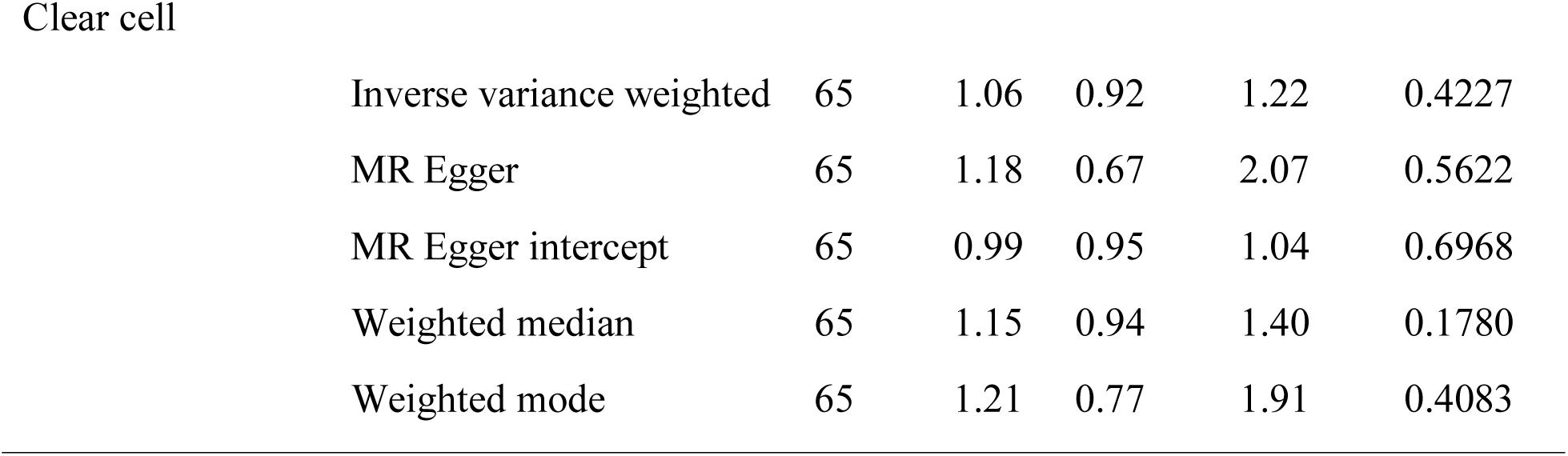
MR of SZ on overall EOC and four ovarian cancer subtypes

|  |  | N |  | Lower | Upper |  |
| --- | --- | --- | --- | --- | --- | --- |
|  | Method | SNPs | OR | 95% CI | 95% CI | *P-value* |
| Overall EOC |  |  |  |  |  |  |
|  | Inverse variance weighted | 66 | 1.09 | 1.04 | 1.14 | 0.0004 |
|  | MR Egger | 66 | 1.18 | 1.00 | 1.39 | 0.0600 |
|  | MR Egger intercept | 66 | 0.99 | 0.98 | 1.01 | 0.3345 |
|  | Weighted median | 66 | 1.10 | 1.03 | 1.18 | 0.0077 |
|  | Weighted mode | 66 | 1.08 | 0.91 | 1.27 | 0.3778 |
| High grade serous |  |  |  |  |  |  |
|  | Inverse variance weighted | 62 | 1.07 | 1.01 | 1.13 | 0.0333 |
|  | MR Egger | 62 | 1.08 | 0.85 | 1.35 | 0.5386 |
|  | MR Egger intercept | 62 | 1.00 | 0.98 | 1.02 | 0.9344 |
|  | Weighted median | 62 | 1.05 | 0.97 | 1.14 | 0.2171 |
|  | Weighted mode | 62 | 1.06 | 0.86 | 1.30 | 0.5857 |
| Mucinous |  |  |  |  |  |  |
|  | Inverse variance weighted | 65 | 1.09 | 0.98 | 1.21 | 0.1258 |
|  | MR Egger regression | 65 | 1.22 | 0.84 | 1.77 | 0.3027 |
|  | MR Egger intercept | 65 | 0.99 | 0.96 | 1.02 | 0.5282 |
|  | Weighted median | 65 | 1.17 | 1.00 | 1.36 | 0.0510 |
|  | Weighted mode | 65 | 1.36 | 0.97 | 1.91 | 0.0797 |
| Endometrioid |  |  |  |  |  |  |
|  | Inverse variance weighted | 64 | 1.10 | 0.99 | 1.22 | 0.0804 |
|  | MR Egger regression | 64 | 1.33 | 0.88 | 1.99 | 0.1778 |
|  | MR Egger intercept | 64 | 0.98 | 0.95 | 1.02 | 0.3478 |
|  | Weighted median | 64 | 1.14 | 0.99 | 1.32 | 0.0753 |
|  | Weighted mode | 64 | 1.23 | 0.87 | 1.74 | 0.2503 |

Clear cell
|  |  |  |  |  |  |
| --- | --- | --- | --- | --- | --- |
| Inverse variance weighted | 65 | 1.06 | 0.92 | 1.22 | 0.4227 |
| MR Egger | 65 | 1.18 | 0.67 | 2.07 | 0.5622 |
| MR Egger intercept | 65 | 0.99 | 0.95 | 1.04 | 0.6968 |
| Weighted median | 65 | 1.15 | 0.94 | 1.40 | 0.1780 |
| Weighted mode | 65 | 1.21 | 0.77 | 1.91 | 0.4083 |

**Figure 1.**
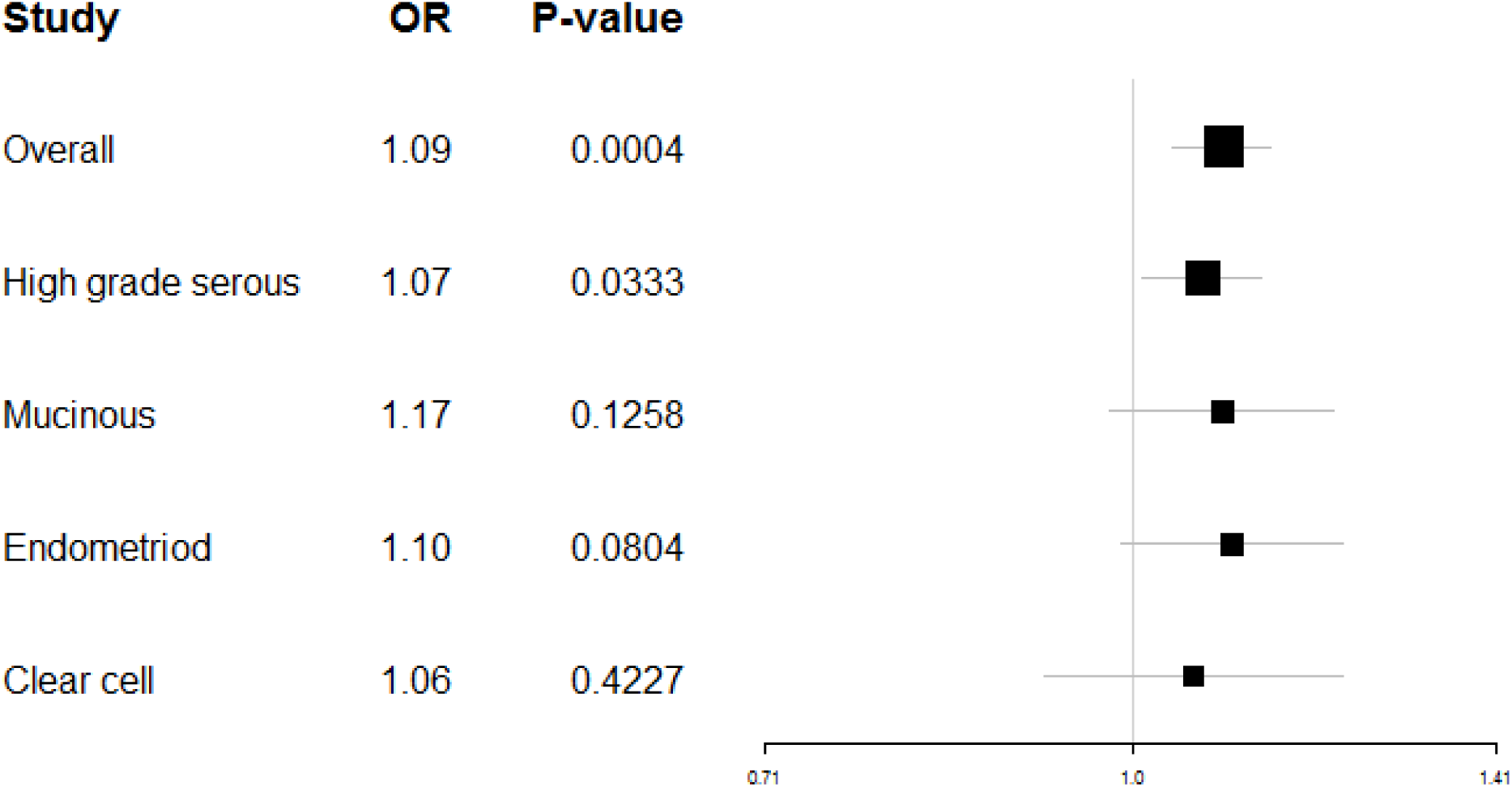
MR results of genetic liability to SZ on overall EOC and four EOC subtypes

Null, increased risks were observed for the remaining subtypes, and IVW radial regression revealed no evidence for heterogeneity: mucinous carcinoma (IVW OR 1.09; 95% CI 0.98, 1.21; *P*=0.1258; *P*[het]=0.7236); endometrioid carcinoma (OR 1.10; 95% CI 0.99, 1.22; *P*=0.0804; *P*[het]=0.9568); and clear-cell carcinoma (OR 1.06; 95% CI 0.92, 1.22; *P*=0.4227; *P*[het]=0.8990). The sensitivity estimators were relatively consistent for all tests (same direction of effect and comparable magnitudes of effect).

### EOC on SZ

Genetic liability to EOC did not causally contribute to risk for SZ (Table 2).

**Table 2.** Results of MR of EOC on SZ.

|  | N |  | Lower | Upper |  |
| --- | --- | --- | --- | --- | --- |
| Method | SNPs | OR | 95% CI | 95% CI | *P-value* |
| Overall EOC |  |  |  |  |  |
| Inverse variance weighted | 9 | 0.95 | 0.87 | 1.04 | 0.2586 |
| MR Egger | 9 | 0.97 | 0.77 | 1.22 | 0.8220 |
| MR Egger intercept | 9 | 0.99 | 0.98 | 1.01 | 0.3345 |
| Weighted median | 9 | 0.94 | 0.86 | 1.03 | 0.2075 |
| Weighted mode | 9 | 0.95 | 0.84 | 1.06 | 0.3791 |

### SZ on GPCs

Two of the 15 tests of SZ on circulating GPCs showed evidence for causal association and 3 had suggestive associations (Table 3). SZ was associated with 1-palmitoleoylglycerophosphocholine (IVW beta log10 increase per log odds higher liability to SZ: −0.021, 95% CI: −0.032, −0.009; *P*=0.0003; *P*[het]=0.7337). Similarly, SZ was associated with 1-oleoylglycerophosphocholine (IVW beta: −0.017, 95% CI: −0.027, −0.007; *P*=0.0006; *P*[het]=0.8507). Suggestive associations were observed for 1-heptadecanoylglycerophosphocholine (IVW beta: −0.019, 95% CI: −0.033, - 0.005; *P*=0.0066; *P*[het]=0.8664); 2-oleoylglycerophosphocholine (IVW beta: −0.014, 95% CI: - 0.024, −0.004; *P*=0.0074; *P*[het]=0.8664); and1-myristoylglycerophosphocholine (IVW beta: - 0.018, 95% CI: −0.031, −0.005; *P*=0.0066; *P*[het]=0.9419). The comparison of the MR estimators showed little evidence for pleiotropy for the GPC metabolites after Bonferroni correction. The suggestively associated GPC metabolites were also robust to the MR sensitivity analyses, except for 1-heptadecanoylglycerophosphocholine for which there was some discrepancy in the direction of effects for the MR estimators, indicating potential pleiotropy. For all the GPCs, including the 10 that had null associations, SZ (Figure 2) lowered the GPCs.

**Table 3.** Results of MR of SZ on circulating glycerophosphocholines (GPCs).

|  | N |  |  | Lower | Upper |  |
| --- | --- | --- | --- | --- | --- | --- |
| Method | SNPs | Beta | Std. Error | 95% CI | 95% CI | P-value |
| 1-palmitoleoylglycerophosphocholine |  |  |  |  |  |  |
| Inverse variance weighted | 60 | -0.021 | 0.006 | -0.032 | -0.009 | 0.0003 |
| MR Egger regression | 60 | -0.032 | 0.026 | -0.082 | 0.019 | 0.2275 |
| MR Egger intercept | 60 | 0.001 | 0.002 | -0.003 | 0.005 | 0.6696 |
| Weighted mode | 60 | -0.049 | 0.020 | -0.088 | -0.009 | 0.0188 |
| Weighted median | 60 | -0.028 | 0.009 | -0.045 | -0.011 | 0.0014 |
| 1-heptadecanoylglycerophosphocholine |  |  |  |  |  |  |
| Inverse variance weighted | 58 | -0.019 | 0.007 | -0.033 | -0.005 | 0.0094 |
| MR Egger regression | 58 | 0.030 | 0.035 | -0.037 | 0.098 | 0.3836 |
| MR Egger intercept | 58 | -0.004 | 0.003 | -0.009 | 0.001 | 0.1516 |
| Weighted mode | 58 | 0.001 | 0.026 | -0.050 | 0.052 | 0.9675 |
| Weighted median | 58 | -0.012 | 0.010 | -0.033 | 0.008 | 0.2298 |
| 1-oleoylglycerophosphocholine |  |  |  |  |  |  |
| Inverse variance weighted | 56 | -0.017 | 0.005 | -0.027 | -0.007 | 0.0006 |
| MR Egger regression | 56 | -0.030 | 0.022 | -0.073 | 0.012 | 0.1694 |
| MR Egger intercept | 56 | 0.001 | 0.002 | -0.002 | 0.004 | 0.5295 |
| Weighted mode | 56 | -0.023 | 0.015 | -0.052 | 0.006 | 0.1314 |
| Weighted median | 56 | -0.019 | 0.007 | -0.032 | -0.005 | 0.0076 |
| 2-oleoylglycerophosphocholine |  |  |  |  |  |  |
| Inverse variance weighted | 57 | -0.014 | 0.005 | -0.024 | -0.004 | 0.0074 |
| MR Egger regression | 57 | -0.016 | 0.024 | -0.063 | 0.031 | 0.5086 |
| MR Egger intercept | 57 | 0.000 | 0.002 | -0.003 | 0.004 | 0.9233 |
| Weighted mode | 57 | -0.006 | 0.015 | -0.036 | 0.024 | 0.7033 |
| Weighted median | 57 | -0.011 | 0.007 | -0.026 | 0.003 | 0.1157 |
| 1-myristoylglycerophosphocholine |  |  |  |  |  |  |
| Inverse variance weighted | 59 | -0.018 | 0.007 | -0.031 | -0.005 | 0.0066 |
| MR Egger regression | 59 | -0.012 | 0.030 | -0.071 | 0.047 | 0.6953 |
| MR Egger intercept | 59 | 0.000 | 0.002 | -0.005 | 0.004 | 0.8347 |
| Weighted mode | 59 | -0.024 | 0.021 | -0.066 | 0.018 | 0.2648 |
| Weighted median | 59 | -0.020 | 0.009 | -0.039 | -0.002 | 0.0332 |

**Figure 2.**
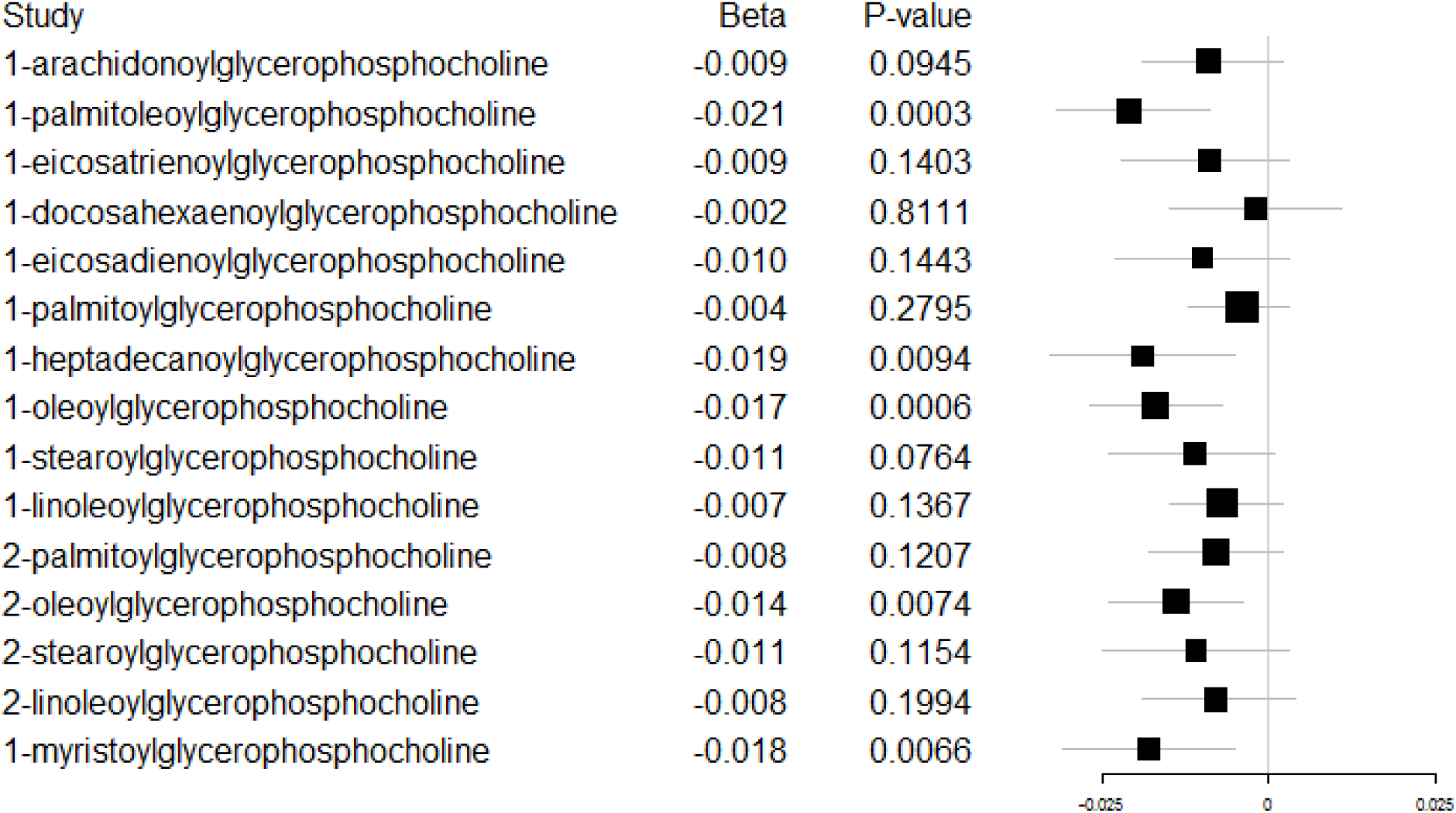
MR results of genetic liability to SZ on 15 GPCs.

The results for the IVW estimates of individual SNPs for all models and the table with the complete list of findings for the MR of SZ on the 15 GPC metabolites, including the null associations, are available in the Supplement.

## DISCUSSION

As the prevalence of SZ ranges from 4 to 7 per 1000 persons worldwide [23], both it and EOC are rare enough that there are few epidemiologic studies examining SZ as a risk factor for EOC. Those that have examined the influence of SZ on EOC have observed mixed findings. Comparing the incidence rate ratio (IRR) of EOC among those with and without SZ, a null but decreased risk for EOC has been reported (IRR 0.98; 95% confidence interval (CI) not specifically provided, but included 1) [24]. A large retrospective study in Sweden of 59,233 patients with SZ reported a null but increased risk for EOC (standardized incidence ratio, SIR, 1.13; 95% CI 0.95, −1.34) among schizophrenics after diagnosis, but a decreased risk for EOC prior to diagnosis (SIR 0.48; 95% CI 0.35, −0.65) [25]. These previous reported associations between SZ and EOC could reflect residual confounding, reverse causation, or measurement error.

A major strength of our investigation is that we leveraged large GWAS to examine these two rare diseases. Our approach optimized power to find associations. A weakness of our study is that we were not able to assess the strength of our MR instruments (F-statistics), due to missing effect-allele frequencies in the Ripke *et al.* (2014) GWAS. However, this is balanced out by the advantage of the two-sample MR design: weak instruments bias causal associations towards the null in two-sample MR. Thus, if our SZ instruments were weak, we likely would not have observed the causal associations we did, as our findings would have been attenuated to the null. Although the MR of SZ on overall EOC was statistically well powered, our analysis of SZ on the EOC subtypes may have been unpowered due to the small number of cases. Thus, we cannot rule out the possibility that SZ is associated with mucinous, clear cell, and/or endometrioid carcinomas, as it was with overall EOC and high-grade serous carcinoma, the most common of the subtypes.

Aberrant choline metabolism is a feature of malignant transformation in breast and ovarian cancer cell lines, where it is marked by a change from low phosphocholine (PCho) and high GPC levels to high PCho and low GPC levels[10], [26], [27]. Aberrant choline metabolism has also been noted clinically for breast cancer patients, with higher GPC levels being implicated in better prognosis [28]. In studies of schizophrenics, high choline concentrations have been observed in patients prior to antipsychotic treatment [29], pointing to perturbation of choline metabolism being independent from antipsychotic treatment and not a result of it. Moreover, GPCs, specifically, have been observed to be statistically lower in patients compared to controls in certain brain regions (antipsychotic use did not appear to be the cause)[30]. Here, we observed that SZ was associated with lower levels of several circulating GPC metabolites and that SZ conferred a weak but notable increased risk for EOC.

Ideally, we would have tested whether these SZ-associated GPC metabolites caused EOC, but we did not have appropriate genetic instruments for this purpose. Instead, we postulate that disturbed choline metabolism may explain the causal association between SZ and EOC. It is also possible that something else may underlie the connection between the two conditions. For instance, Brown (2016) proposed that SZ may involve cell transformation, similar to that of early stage cancer initiation or attenuated tumorigenesis [31] -- but without taking on a complete tumor phenotype. This is a provocative hypothesis that fits with our observations but may also, if true, reflect other early features of cancer, perhaps immune rather than metabolic. Another possibility is that changes in choline metabolism in SZ perturb the one-carbon cycle. While free choline is primarily used in phospholipid synthesis, excess choline serves as a source of one-carbon units for methylation; choline can directly re-methylate homocysteine to contribute to methionine formation and homeostasis [32]. Thus, perhaps SZ-induced aberrant choline metabolism mimics the “cholinic phenotype” observed in EOC and/or contributes to it, maybe via perturbation of the one-carbon cycle.

Minimally, our results highlight the possibility that something about SZ imparts risk for developing EOC. Understanding this risk has implications for the biology of both diseases and could aid efforts at earlier detection of EOC, especially if the biological process perturbed in the two conditions also occurs in the absence of SZ. Investigating alterations in choline metabolism is a starting point to focus future investigation.

## ACKNOWLEDGEMENTS

We have no acknowledgements.

